# Single-cell transcriptomics to define *Plasmodium falciparum* stage-transition in the mosquito midgut

**DOI:** 10.1101/2022.04.05.487115

**Authors:** Mubasher Mohammed, Alexis Dziedziech, Vaishnovi Sekar, Medard Ernest, Thiago Luiz Alves E Silva, Balu Balan, S. Noushin Emami, Inna Biryukova, Marc R. Friedländer, Aaron Jex, Marcelo Jacobs Lorena, Johan Henriksson, Joel Vega Rodriguez, Johan Ankarklev

**Affiliations:** Department of Molecular Biosciences the Wenner Gren Institute, Stockholm University, Svante Arrhenius väg 20C, SE-106 91, Stockholm, Sweden; Science for Life Laboratory, Department of Molecular Biosciences, The Wenner-Gren Institute, Stockholm University, Svante Arrhenius väg 20C, SE-106 91, Stockholm, Sweden; Laboratory of Malaria and Vector Research, National Institute of Allergy and Infectious Diseases, National Institutes of Health, 12735 Twinbrook Parkway, Rockville, MD 20852, USA; Population Health and Immunity Division, The Walter and Eliza Hall Institute of Medical Research, Parkville, Melbourne, VIC, Australia; Faculty of Veterinary and Agricultural Sciences, The University of Melbourne, Parkville, VIC, Australia; Department of Molecular Microbiology and immunology, Bloomberg School of Public Health, Johns Hopkins University, 615 N Wolfe St, Baltimore, MD 21205, USA; Laboratory for Molecular Infection Medicine Sweden (MIMS), Department of Molecular Biology, Umeå University, SE-901 87, Umeå, Sweden; Microbial Single Cell Genomics, Department of Cell and Molecular Biology and Science for Life Laboratory, Uppsala University, Uppsala, Sweden

## Abstract

Malaria inflicts the highest rate of morbidity and mortality among the vector-borne diseases. The dramatic bottleneck of parasite numbers that occurs in the gut of the obligatory vector mosquito provides a promising target for novel control strategies. Using single-cell transcriptomics, we analyzed *Plasmodium falciparum* development in the mosquito gut, from unfertilized female gametes through the first 20 hours post blood feeding, including the zygote and ookinete stages. This study revealed the transcriptional trajectories of the ApiAP2 family of transcription factors, and of parasite stress genes in response to the harsh environment of the mosquito midgut. Further, employing structure-based functional predictions we found several upregulated genes predicted to encode intrinsically disordered proteins (IDPs), a category of proteins known for their importance in regulation of transcription, translation and protein-protein interactions. IDPs are known for their antigenic properties and may serve as suitable targets for antibody or peptide-based transmission suppression strategies.

## Background

Malaria remains a large global health burden, infecting approximately 224 million people each year and having a death toll exceeding 627,000, mainly children in Sub-Saharan Africa (*World Malaria Report 2021*, n.d.). Out of the five *Plasmodium* species known to infect humans, *P. falciparum* causes the vast majority of severe cases and deaths. *Plasmodium* parasites have a complex life cycle that extends between the human host and the female *Anopheles* mosquito. Parasite development within the mosquito begins during the ingestion of a blood meal containing male and female gametocytes. Gametocytes quickly differentiate into gametes in the mosquito midgut, which fuse to form a diploid zygote. The zygote differentiates into a motile banana-shaped ookinete that migrates within the blood bolus, traverses the midgut epithelium, and forms an oocyst on the basal side of the midgut. During the transition from the blood bolus to the hemocoel, the parasite is exposed to harsh conditions within the midgut, including immune factors from the human host, the midgut microbiome and oxidative stress from blood digestion (Billker et al., 1997; Smith et al., 2014). Subsequently, the ookinetes lodge into the basal lamina, where they develop into replicative oocysts and thousands of sporozoites are produced. Sporozoites are released from mature oocysts into the hemocoel, from where they invade the salivary glands and are delivered into a vertebrate host when the mosquito takes a blood meal (Graumans et al., 2020).

While *P. falciparum* undergoes large scale asexual replication in the human host, sexual recombination occurs only in the mosquito vector. This enables genetic crossing and spread of genetic factors such as drug tolerance, into the parasite progeny (Vaughan et al., 2015). The human-to-mosquito transmission phase is one of the major bottlenecks in the parasite’s lifecycle, in part due to the limited number of gametocytes taken up by the mosquito, but also due to immune factors from the human blood, the mosquito microbiome and the effective innate immune response elicited by the mosquito (Smith et al., 2014). Moreover, the anopheline mosquito has evolved to express a battery of antimicrobial defenses including long non-coding RNAs, nitric oxide, prophenoloxidases, and anti-microbial peptides (Dimopoulos et al., 2001; Padrón et al., 2014). These events contribute to the large bottleneck in parasite numbers during early development in the mosquito midgut, making these stages a crucial target for transmission-blocking interventions (Griffin et al., 2010). In addition, recent advances in malaria control have bolstered the interest in malaria intervention strategies linked to the reduction of transmission, including vaccines that target sexual, sporogenic, and/or mosquito stage antigens in order to interrupt malaria transmission (SSM-VIMT) (Sauerwein & Bousema, 2015). Although *P. falciparum* development in the mosquito midgut is considered a prime target for the development of effective transmission-blocking interventions, little is currently known about the transcriptional program that controlling these processes, with surprisingly few proteins characterized and annotated. In recent years, single-cell RNA sequencing (scRNA-seq) has been used extensively to resolve cellular heterogeneity, discern key transcriptionally regulated biological processes and predict transcript and protein expression patterns and functions in multiple stages during the life cycle of *P. falciparum* (Howick et al., 2019; Ngara et al., 2018; Poran et al., 2017; Real et al., 2021; Reid et al., 2018). Nonetheless, a detailed and comprehensive transcriptional map of key *P. falciparum* developmental stages in the mosquito midgut is still missing (Figure 1A).

**Figure 1.**
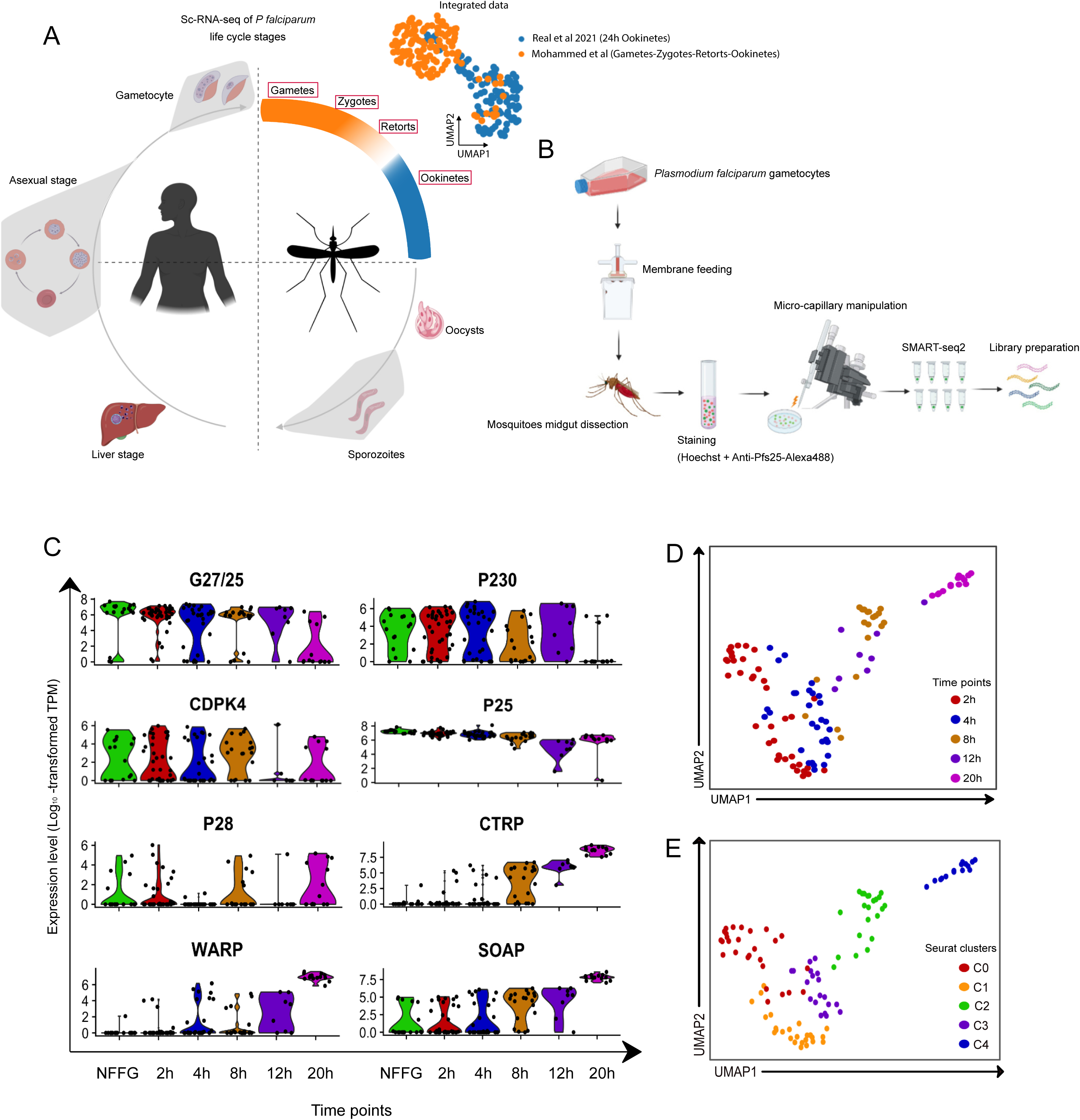
Transcriptomic analysis of single *P. falciparum* parasites isolated from the midgut blood bolus of *An. gambiae* mosquitoes. **A**. Schematic diagram of the *P. falciparum* lifecycle, highlighting the coverage of single-cell transcriptome data across the lifecycle that makes up the current malaria atlas. The UMAP shows an overlay of the scRNA-seq data of 24h ookinetes from Real *et al*., 2021 (blue) and the current study (orange, top right) (Illustrations created using www.biorender.com). **B.** Schematic diagram of the experimental pipeline: *in vitro* cultured gametocytes were fed to *An. gambiae* female mosquitoes by standard membrane feeding. Mosquito midguts were dissected at six different time points, homogenized, stained and the homogenate was placed in an inverted fluorescence microscope. Parasites were selected based on their positivity to Hoechst and Anti-Pfs25-Alexa-488 staining by fluorescence microscopy and isolated by micro-manipulation. cDNA libraries were prepared for each cell using a modified version of the Smart-seq2 protocol. **C**. Violin plots of the average expression level (Log_10_ transformed TPM) of marker genes for *P. falciparum* midgut stages. The marker genes were selected from Bennink *et al*. 2016 for validation of our single-cell transcriptome dataset. **D** and **E**. UMAP of the single-cell transcriptome data overlaid with the collection time points at which they were isolated (D) or the respective Seurat clusters (E). These plots show the two-dimensional projection of five isolated time points. The global transcriptome similarities and differences were assessed using a k-nearest neighbors (kNN) force-directed graph on the first 10 Principal Components with true signal variation from our single-cell transcriptome dataset based on Elbow plot.

In this study, we use scRNA-seq to explore the *P. falciparum* developmental dynamics from unfertilized female gametes, through the zygote and the ookinete stages. All parasite cells were carefully isolated by micromanipulation from infected *Anopheles gambiae* mosquitoes. We sought to elucidate the timing of expression of genes connected to cytoskeletal modifications, invasion, and meiosis. Computational analyses identified the timing of expression for members of the ApiAP2 family of transcription factors throughout the developmental timeline. Further, expression of a set of genes containing the upstream binding motif of the stage-specific ApiAP2-O were connected to the expression of this transcription factor. Moreover, we provide insights into the connections of the parasite’s stress responses during the mosquito midgut stage, with morphogenic changes and evolution of cell fate. Finally, we identify several highly expressed genes with non-annotated function, that are predicted to be intrinsically disordered proteins (IDPs) and novel immunogenic candidates for antibody-based therapies. The elucidation of genetic factors involved in *P. falciparum*’s differentiation in the mosquito midgut provides vital insights into the biology of these vulnerable developmental stages while identifying new targets for the control of malaria transmission.

## Results

### Unsupervised clustering defines a developmental trajectory and transcriptional heterogeneity among the early developmental timepoints

To enable high-resolution transcriptional profiling of *P. falciparum* as it develops in the lumen of the mosquito midgut, we infected the African main malaria vector *An. gambiae* with mature stage V gametocytes of the NF54 genetic background. Mosquito midguts were dissected at 2, 4, 8, 12, and 20 h post infection (p.i.), homogenized to release the blood bolus content, followed by Hoechst (DNA) and anti-Pf25 antibody staining to label the parasite sexual stages (female gametes, zygotes and ookinetes). Parasites do not develop synchronously in the midgut of the mosquito and therefore, we sought to visualize parasites prior to isolation through DNA and Pfs25 staining, and select only parasites with intact morphology. Individual parasites were collected by micro-capillary manipulation within 30 minutes post dissection and homogenization (Figure 1B), and cDNA libraries were prepared for each isolated parasite using a modified version of the Smart-seq2 protocol (Ngara et al., 2018; Reid et al., 2018) using 24 cycles for cDNA amplification.

A total of 180 single cells were collected, of which 125 single cells were retained for subsequent analyses after filtering out cells with poor read counts. The remaining cells were quality controlled based on mRNA count and number of genes expressed (Supplementary Figure 1A). The data from each time point were validated by analyzing the expression levels of known marker genes for specific midgut developmental stages, including G27/25, p230, CDPK4, P25 and P28 (Bennink et al., 2016). Marker genes associated with ookinete development or midgut invasion like CTRP, WARP and SOAP, showed significantly elevated expression levels towards the end of the time course (Figure 1C). Further, we integrated and normalized our data with a recent single-cell data set published by Real and colleagues (Real et al., 2021) in order to validate the coverage of the hitherto unexplored *Plasmodium* developmental stages. As seen in Figure 1A, our late, 20h timepoint aligns with the majority of cells from the Real *et al*. dataset, which were collected at 24 h post mosquito infection, while our earlier timepoints form separate clusters (Figure 1A and Supplementary Figure 1B). A total of 2,000 genes with significant variance were selected for downstream analyses, while 2,835 did not show significant variance (Supplementary Figure 1C). Further, among the genes with the highest average change, we found several non-annotated genes and genes that were previously shown to have essential roles during parasite development in the midgut, such as GAMER, PIMMS1, PIMMS57 and SOPT (Supplementary Figure 1D) (Akinosoglou et al., 2015; Ukegbu et al., 2020). Next, we clustered the cells in an unsupervised manner using UMAP non-linear dimensionality reduction, which identified five distinct clusters that largely correspond to the time points at which parasites were collected (Figure 1D and E, see Methods for details). The UMAP displays transcriptional variability between and within isolated time points. Early timepoints, in particular, were computationally predicted to diverge by cell type within each time point (Supplementary Figure 2A).

### Pseudotime alignment indicates heterogeneity among the early timepoints

Given the heterogeneity of the identified cells, we sought to determine a biologically relevant temporality to the cell clusters. Since the non-fertilized female gametes (NFFGs) were cultivated *in vitro* and therefore not exposed to stresses from the mosquito midgut, we performed a cell trajectory pseudotime analysis that included non-fertilized female gametes. The results indicated that the inclusion of these cells created a second trajectory, which appeared to distort the natural developmental trajectory as visualized in (Supplementary Figure 2B). Therefore, we excluded the 0- hour (*in vitro*) time point from subsequent pseudotime analyses. To infer a pseudotime alignment for the five clusters identified by Seurat analyses, we reconstructed the cell trajectory in terms of their relative developmental stage. Clusters were reordered with C3 indicated to be the earliest developmental stage, followed by C1, C0, C2 and ending with C4 as the terminal cluster (Figure 2A). Further, when the data was ordered along a temporal plane, i.e., the pseudotime (pt) axis, three distinct clusters, pt0, pt1 and pt2 (Figure 2A, right) were delineated. The pt lineage showed general similarity with the sampling timepoints, although the early timepoints showed a trend of asynchronicity where some of the cells aligned with a more progressed stage of development while others aligned with a differing transcriptional state, which we hypothesize may indicate stalled development (Figure 2B, left). Further, the Seurat clusters showed greater spread over the pseudotime within a single group and more overlap between groups in comparison to the pt clusters, which were more distinct (Figure 2B, middle *vs*. right). While the pt clusters showed more mutual exclusivity than Seurat clustering, the Seurat clustering indicated more distinct intermediary points in the transition from the early zygote to the ookinete. We utilized the Seurat cluster variance when exploring the role of key genes in transcriptional programing while we relied on slingshot pseudotime clusters for determining distinct biological processes. The GO enrichment for the pt clusters revealed similar GO terms to the Seurat clustering, pointing to an ordered and predictable transcriptional regulation of gene expression programs. For example, pt0 was related to “response to stimulus”, “localization” and “reproduction”, pt1 was related to “DNA replication”, “signaling”, and “translation”, while pt2 was related to “entry into host”, “invasion”, and “response to abiotic stimulus” (Figure 2C). Based on the imaging data acquired during the isolation of individual parasites, we saw distinct cell morphologies throughout the time of collection (Figure 2D). When comparing the nucleus mean volume, fluorescence intensity and mRNA abundance, we saw variation across the 5 Seurat clusters with C1 containing higher nuclear mean volume and fluorescence intensity, which coincides with the post meiotic tetraploidy (4N) (Supplementary Figure 3 A, B and C). To identify the genetic signature for each cluster, we generated the scaled average expression of the top10 differentially expressed genes for the Seurat clusters (C0-C4) and the top 20 differentially expressed genes for the pt clusters (pt0-2) (Supplementary Figure 4 A and B, respectively). Thus, based on morphology, nuclear DNA and GO term analyses, clusters C3 and C1 align with pt0 and represent early zygote development, C0 aligns with pt1 and represents the maturing zygote to early ookinete development, whereas C2 and C4 align with pt2 and represent mid-to-late ookinete development.

**Figure 2.**
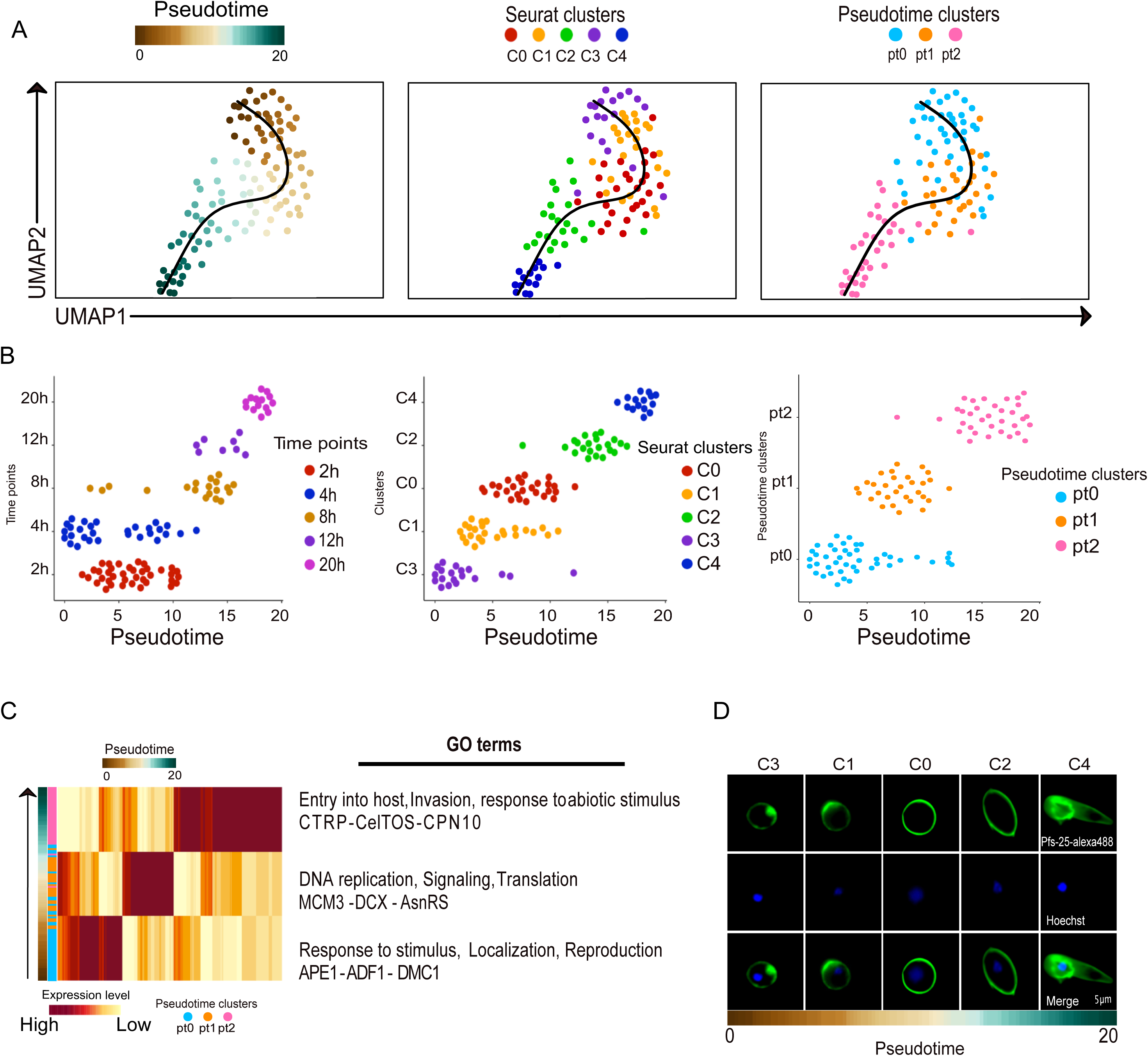
Ordering cells based on the developmental trajectory. **A**. Ordering of cells along their pseudotime developmental trajectory using Slingshot. Left Panel: Pseudotime alignment of cells used in this study, where each dot represents individual parasites and their positioning is determined by the total relative expression in comparison to the other parasites included in the analysis. The color coding represents their predicted development along the pseudotime axis. Middle panel: The pseudotime developmental trajectory overlaid with the assigned Seurat clusters, indicating that C3 is at the root of the trajectory and C4 represents the terminal state. Right panel: The pseudotime line overlaid with three unique mRNA patterns based on overarching similarity among major cell communities formed during the process of lineage reconstruction. **B**. Slingshot ordering of cells plotted over a pseudotime axis. Left panel: cells from each collection timepoint plotted on the pseudotime axis. Middle panel: cells representing each of the five Seurat clusters plotted on the pseudotime axis. Right panel: pseudotime clusters plotted on the pseudotime axis. **C**. Heatmap showing the average transcript abundance across the pseudotime (pt) axis, based on the Slingshot clustering (pt0, pt1 and pt2). Notable GO terms are indicated for each of the three pt clusters. **D**. Representative Z-stack images of developmental stages based on the ordering of the Seurat clusters (C0-C4) over the pseudotime axis. Images depict the surface marker P25-Alexa488 (green, top panel), nuclear Hoechst stain (blue, middle panel), and merged P25-Alexa488+Hoechst images (bottom panel).

### Pseudotime ordering identifies gene programs that drive *P. falciparum* development in the midgut

To gain insight into the transcriptional programing for the clusters identified over the pseudotime, we performed hierarchical clustering and gene ontology analysis of the top 200 differentially expressed genes across all timepoints (Figure 3A, Supplementary Table 1). We observed a sequence of biological events corresponding to the development from early zygote to ookinetes over the pseudotime, as supported by the top scoring GO terms which included “DNA Replication” in pt0, “Reproduction” in pt1 and “Entry into host” and “Negative regulation of metabolic process” in pt2. After performing a gene set enrichment analysis over the pseudotime, increasingly fewer cells showed enrichment for metabolic processes. The average expression of each significantly upregulated program was assessed throughout the developmental time course (Supplementary Figure 5 A and Supplementary Table 2). To further define patterns of biological processes among the gene modules that make up the pseudotime clusters, we also ran over-representation analyses of the pt clusters’ (pt0 to pt2) gene ontology terms (Supplementary Figure 5 B, C and D).

**Figure 3.**
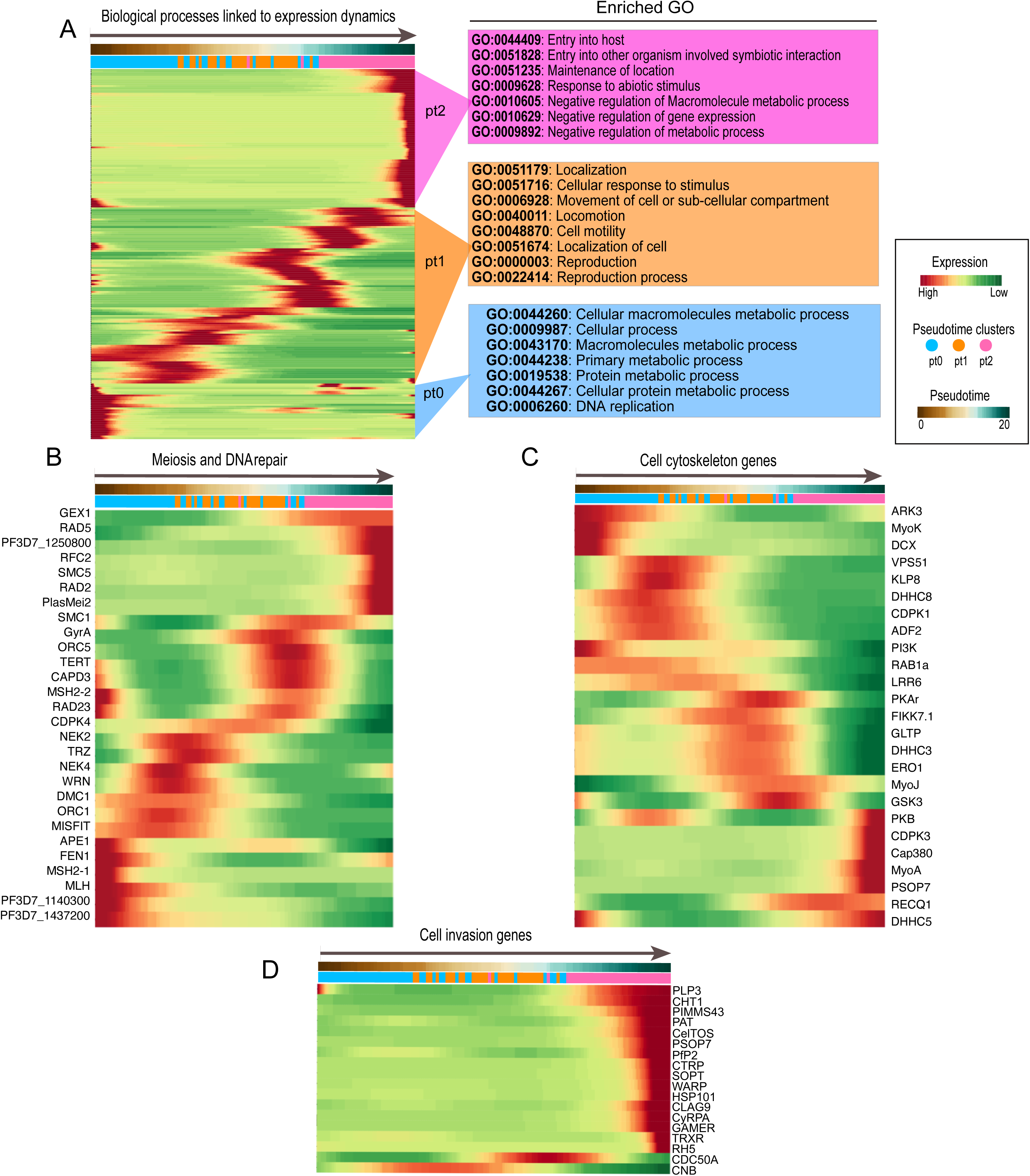
Top differentially expressed genes across the pseudotime axis. **A**. Hierarchical clustering of the top 200 expressed genes across the Slingshot pseudotime axis. The genes were grouped according to their ontology terms and analyzed for the top GO terms corresponding to each pseudotime cluster is indicated to the right of the heatmap. **B-D**. Heatmap of genes directly or indirectly involved in meiosis and DNA replication (B), cytoskeleton remodeling (C) and cell invasion (D) across the Slingshot pseudotime axis. For all heat maps, genes were selected based on their biological profile and annotation in Plasmodb and clustering was based on the Euclidean distance of scaled rows.

Given that fertilization and meiosis occur in the midgut under extreme oxidative stress conditions, we selected known meiosis and DNA repair genes to further our understanding of these processes during the midgut developmental stages (Figure 3B). A distinct set of early expressed genes, including APE1, FEN 1, MSH2-1, MLH, PF3D7_1140300 and PF3D7_1437200, were included in DNA repair and DNA replication. A subsequent set of genes that correspond to developmental progression through meiosis in the early zygote, included the expression of MISFIT (male-inherited sporulation factor important for transmission), FEN1, a DNA repair enzyme, and DMC1, a meiotic recombination protein, and MLH, a DNA mismatch repair protein. MISFIT may also contribute to cell cycle progression and replication dynamics in conjunction with NEK4 and ORC1 leading to the development of the early ookinete (Bushell et al., 2009). A third set of genes expressed at an even more advanced stage of development include: RAD23 involved in DNA repair, GyrA, DNA topoisomerase activity, as well as MSH2-2 involved in DNA recombination, Capd3, meiotic chromosome condensation, TERT, telomere maintenance and SMC1 involved in sister chromatid segregation. In summary, the high-resolution single-cell dataset enables the description of expression timing of key genes involved in meiosis and its coordination. Interestingly, we observe a difference between ORC1 and ORC5 expression, which may indicate how the timing of the different subunits of the origin recognition complex in *P. falciparum* are orchestrated in their expression.

Towards the end of the pseudotime, there are seven genes, including RAD2, RAD5 and PF3D7_1250800, involved in DNA repair that may be expressed in response to oxidative stress caused by heme released from hemoglobin digestion, or alternatively expressed in late ookinetes in preparation for the subsequent mass replication during sporogony. The additional four genes RFC2, involved in DNA replication, SMC5, structural maintenance of chromosomes, GEX1, karyogamy and PlasMei2, linked to schizogony, are likely to function during sporogony in oocysts.

Considering the extensive morphological changes that occur as *P. falciparum* develops from early zygotes to mature ookinetes, we sought to better understand the coordination of its cytoskeletal gene network (Figure 3C). We found that ADF2, MyoK, and DCX, involved in actin depolymerization, cytoskeletal regulation and protein polymerization, respectively, were highly expressed in pt0, as were Pi3k and Rab1a, indicating a potential role for endosome recycling in the early development of zygotes. Pt1 showed upregulation of VPS51 and DHHC8, potentially increasing vesicle sorting and protein transport, followed by the upregulation of DHHC3, which is involved in protein palmitoylation and FIKK7.1, involved in protein phosphorylation and signal transduction. Thus, pt1 likely portrays the coordination of vesicles and protein transport that aid in parasite survival in the gut and development from early to mid/late ookinetes. Finally, cells in pt2 significantly expressed other regulators of actin (ADF2) and cell division (MyoJ) as well as ARP-dependent DNA helicase Q1 (RECQ1), which plays a role in DNA double-strand break repair and DNA unwinding (Claessens et al., 2018), possibly in preparation for the subsequent sporogony. Further, late ookinete proteins like Cap380 (oocyst capsule protein) essential for oocyst formation (Srinivasan et al., 2008) and PSOP7 (putative secreted ookinete protein) linked to midgut epithelial invasion (Ecker et al., 2008), were both highly expressed in preparation of epithelia invasion and subsequent oocyst development (Figure 3C). Thus, our data shines new light on the complex transcriptional orchestration of cytoskeletal changes as zygotes differentiate into ookinetes, and as ookinetes prepare for oocyst formation.

We also analyzed the timing of expression of ookinete invasion genes, which are almost exclusively expressed within pt2. These include well-known ookinete marker genes such as; Gamer, SOPT, WARP and PIMMS43 (Figure 3D). Interestingly, Clag9, a cyto-adherent molecule, known to form complex with RHOPH1 during erythrocyte invasion and nutrient uptake (Schureck et al., 2021), was found to be upregulated in late ookinetes.

### Defining the temporality of expression of ApiAP2 family members

Members of the ApiAP2 transcription factor family, which in *P. falciparum* consists of 27 genes, regulate differentiation and stage progression events (Jeninga et al., 2019). We observed significant upregulation of expression in four out of the five AP2-O genes expressed during parasite development in the mosquito midgut, AP2-O4 did not show expression at any time point (Figure 4A). AP2-O was found to be expressed in the C0 and C1 clusters, indicating high expression in the early zygote and in agreement with a previous report showing AP2-O transcription in stage V female gametocytes (Kaneko et al., 2015). AP2-O2, AP2-O3 and AP2-O5 show expression in C0 and C2 (Figure 4A), during the zygote to ookinete transition. Thus, the timing of expression of the AP2-O genes supports a role in the regulation of genes linked to ookinete function, as previously shown by functional analyses (Li et al., 2021; Modrzynska et al., 2017, p. 2; Zhang et al., 2017). Interestingly, we observe additional AP2 genes that appear to be upregulated in a significant proportion (>50%) of cells within their respective clusters (Figure 4A). These include two additional AP2 genes in C2 (PF3D7_1239200 and PF3D7_1342900) and four AP2 genes in C4 (PF3D7_0420300, PF3D7_0934400, PF3D7_0802100 and PF3D7_1139300), none of which has previously been linked to any developmental stage. In agreement with our data, PF3D7_0934400 was shown to be highly expressed among ookinetes in the single-cell data by Real and colleagues. Also, PF3D7_0622900: AP2Tel, which is significantly upregulated in C4, appears to show low to intermediate expression among ookinetes in the same dataset (Real et al., 2021).

**Figure 4.**
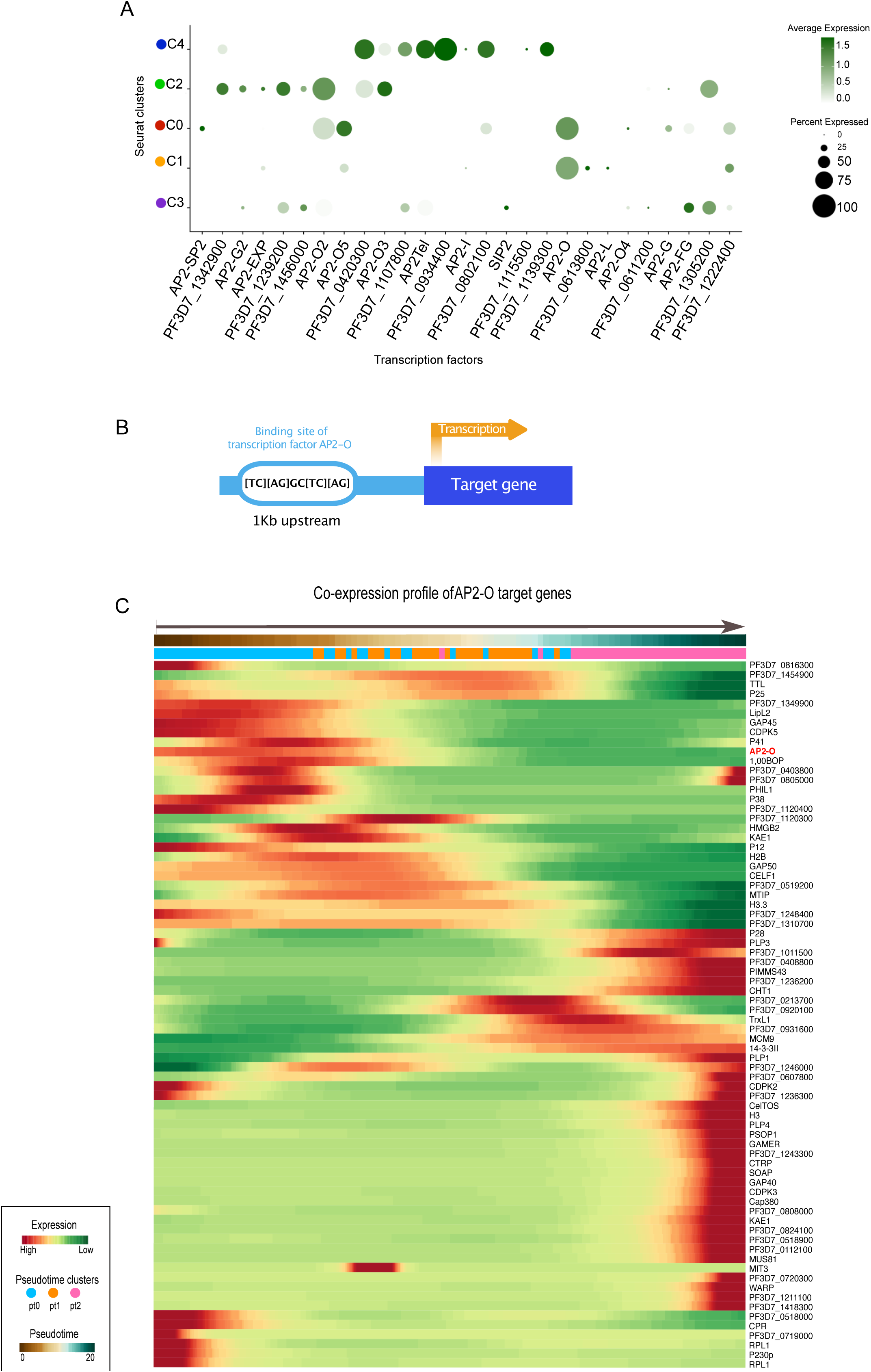
Gene expression patterns of the ApiAP2 transcription factors throughout the *P. falciparum* development in the mosquito midgut. **A**. Dot plot showing the average expression and the proportion of cells that express each of the 27 ApiAP2 transcription factors across the Seurat pseudotime clusters. **B**. Schematic illustration of the process of validating genes harboring an ApiAP2-O binding site, including a motif screen of the 1 KB upstream region from the transcription start site of select genes. **C**. Pseudotime heatmap demonstrating the timing of expression of the ApiAP2-O transcription factor and its putative cis-target genes. Scaled expression values of the genes were ordered based on high (red) or low (green) differential gene expression (DGE) across the pseudotime axis.

To gain further insight into gene regulation during *P. falciparum* zygote to ookinete development, we made use of previously published ChIP-seq data on *P. berghei* parasites to identify genes regulated by AP2-O (Kaneko et al., 2015). The target genes from this study were used to find orthologs in the *P. falciparum* genome that showed significant upregulation in our scRNA-seq data. To validate the orthologs, we investigated the presence of the AP2-O binding motif [TC][AG]GC[TC][AG] (Jeninga et al., 2019; Kaneko et al., 2015) within a 1 kb region upstream of the start codon of differentially expressed genes (Figure 4B). We found that a small number of genes putatively regulated by AP2-O, including *p12*, PF3D7_1349900, PF3D7_0518000 and *cpr*, were upregulated at the beginning of the pseudotime followed by downregulation at an earlier or similar timepoint as the expression of AP2-O (Figure 4C). The *p12* gene belongs to the 6-cys protein family expressed in blood-stage parasites, and was detected in the proteome of merozoites (Sanders et al., 2005). The *cpr* gene has been predicted to encode NADPH-cytochrome P450 reductase, with a predicted role in oxidation-reduction (GO:0055114, AmiGO, geneontology.org). PF3D7_1349900 and PF3D7_0518000, are both unknown proteins but have previously been shown to be transcribed in mature gametocytes and to some level in ookinetes (López-Barragán et al., 2011).

To compare the genes expressed in our scRNA-seq dataset and containing the AP2-O binding motif with the orthologs in the study by Kaneko and colleagues (Kaneko et al., 2015), we only included genes expressed by at least 25% of the cells in each pt cluster. Due to the nature of dropouts in single-cell sequencing data and the conservative threshold set in our analysis, our list of genes is not as extensive as the one by Kaneko et al. (Kaneko et al., 2015). However, our analysis shows a significant overlap with the target genes validated in Kaneko *et al*. We detect significant expression of genes involved in midgut epithelium invasion and oocyst formation, including the secreted proteins; CelTOS, CHT1, CTRP, GAMER, PSOP1, SOAP and WARP, the plasma membrane associated protein; P25, the oocyst development protein Cap380, and microneme proteins; PLP3, PLP4. Furthermore, the pellicular protein TTL, histone proteins H2B, H3 and H3.3, DNA replication proteins MCM9 and CDPK3, which are essential for midgut infection, are all expressed in our dataset. Finally, PIMMS43, previously described as having a function in parasite immune evasion and sporogonic development (Ukegbu et al., 2020) and MUS81 involved in DNA-repair (Lee et al., 2014) were also expressed.

### Response to stress during mosquito midgut development

The stress response elicited by *P. falciparum* is an essential mechanism for adaptation to host environmental factors across the lifecycle (Duran-Bedolla et al., 2017). We investigated the transcriptional activity of known cellular stress marker genes and found transcriptional differences between the Seurat clusters, C0-C4 (Figure 5A). The cells in the C0 cluster showed higher average expression of nPrx, which encodes a nuclear peroxiredoxin that has been suggested to protect nuclear components from oxidative damage or be involved in DNA repair during the asexual blood stage (Richard et al., 2011). Further, genes such as TRXR, PF3D7_0213500, 1-CysPxn and DHX36, are specifically expressed during the later developmental stages C4 (Figure 5A). We also compared the stress genes in non-fertilized female gametes (NFFGs) which showed a smaller population of cells expressing stress related genes compared to C0 (Figure 5A and B). The genes expressed in *in vitro* generated NFFGs, likely include genes expressed through a preprogrammed default stress pathway in comparison to the larger number of heterogeneous genes expressed in C0. The population of cells in C0 likely demonstrate two trajectories for cells either fated to properly maturate into an ookinete or parasites that arrested and failed to mature.

**Figure 5.**
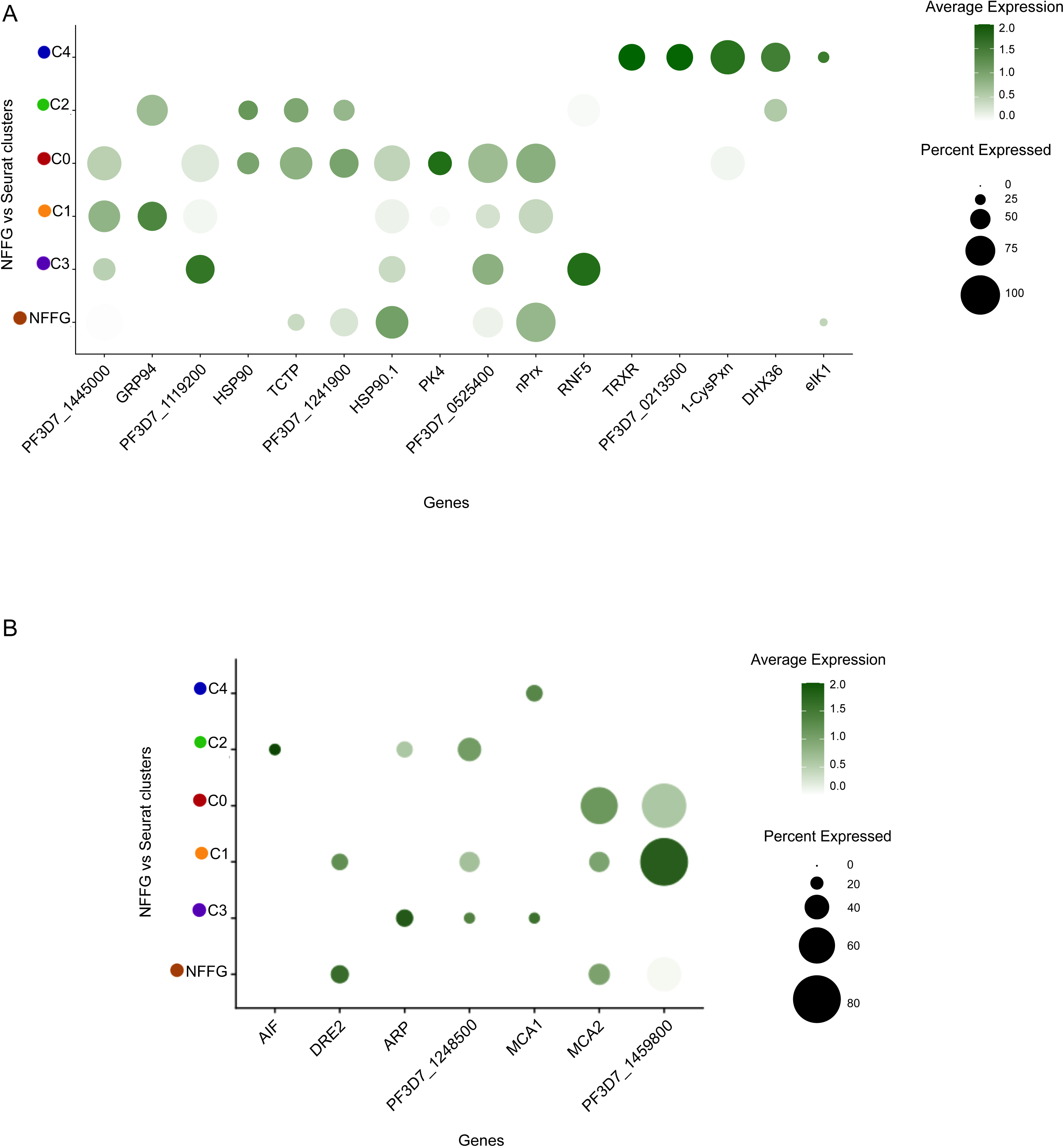
Timing of expression of genes linked to parasite cellular stress and apoptosis. **A**. Dot plot comparing the average expression of stress response genes among cell clusters (post-infection) versus non-fertilized female gametes (NFFG) isolated directly from cell cultures. **B**. Dot plot comparing the average expression of apoptosis-related genes between post-infection cell communities inside the mosquito and NFFGs.

### Protein modelling of stage-specific non-annotated genes

Due to the ability of membrane proteins to induce immune responses as well as their importance in nutrient/chemical uptake (Kirk & Horner, 1995), we sought to define and characterize non-annotated membrane proteins, which make up approximately 33% of the *Plasmodium* genome (Böhme et al., 2019). Such proteins could serve as targets in novel antibody-based therapies. First, from the pt clusters, we identified annotated and non-annotated genes that were significantly upregulated (P adjusted value <0.05) (Figure 6A). From pt1 and pt2, 41 and 111 non-annotated gene candidates made the described cut-off, respectively. The tables in Figures 6B and 6C denote the presence or absence of predicted TM domain and/or a signal peptide. Of note, all the selected non-annotated genes in Figures 6B and 6C are classified as intrinsically disordered proteins (IDPs), all of which had a minimum average log fold expression of 0.5 (Supplementary Table 5). These proteins are characterized by the presence of large segments of disordered structure under normal physiological conditions. Distortion exceeding 15.5% of the entire protein secondary structure were classified as IDPs (Figure 6D, Supplementary Figure 6). IDPs have been described as highly immunogenic (Guy et al., 2015) and may be candidates for novel antibody or peptide-based therapeutics. Molecular simulation (Supplementary Figure 7) shows how protein structure can be predicted for therapeutic molecules. This is an important tool for the identification of immunogenic proteins to be used for vaccine development.

**Figure 6.**
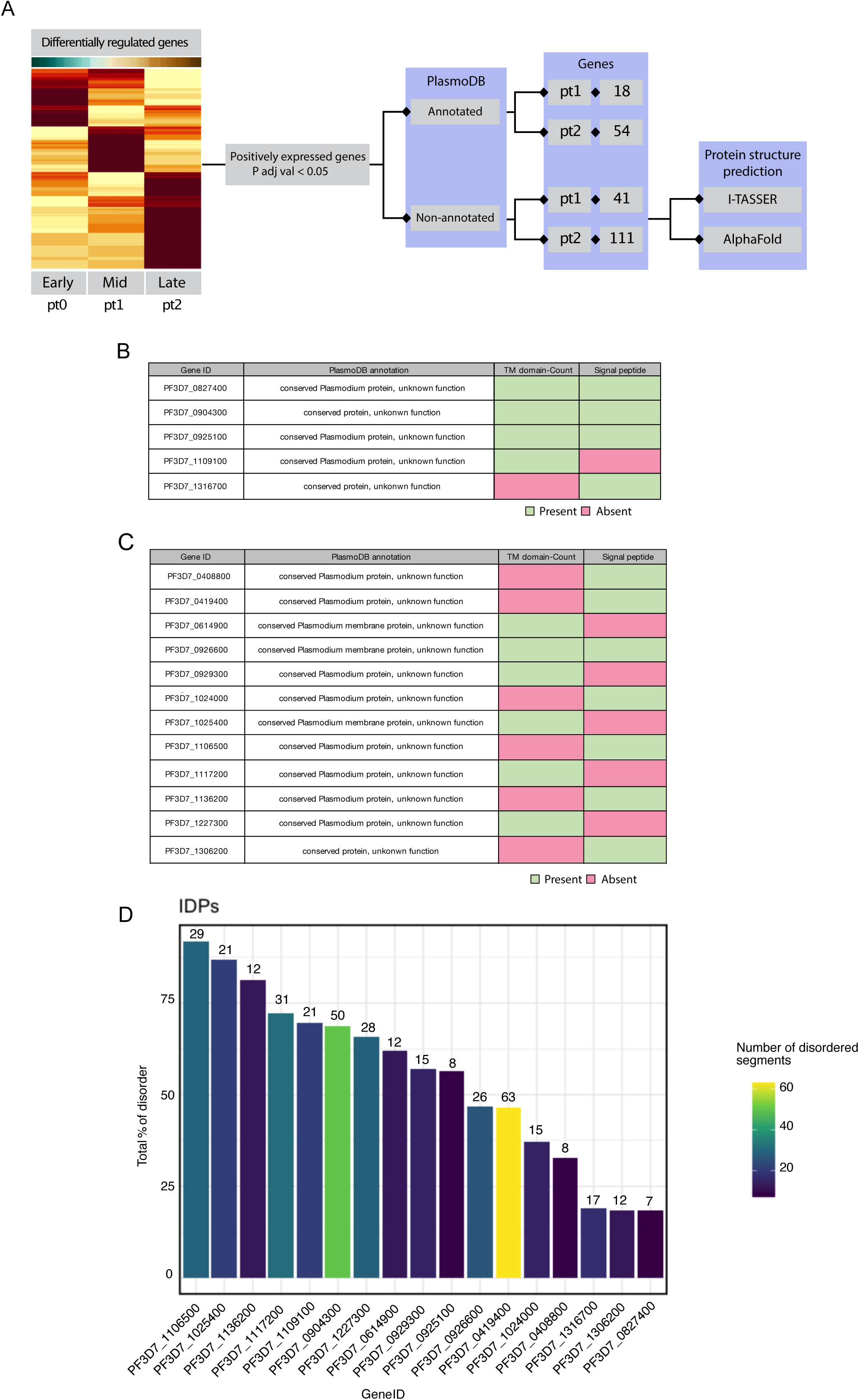
Structural predictions of putative therapeutic targets. **A**. Flow diagram depicting the selection process of highly expressed, non-annotated genes. Genes with *P* adj values of < 0.05 were selected in pt1 and pt2, and further analyzed using PlasmoDB, I-TASSER and Alpha-fold, which indicate a significant likelihood that certain genes encode membrane proteins and signal peptides. **B**. Table of genes from (A) linked to the pt1 cluster, showing the PlasmoDB annotation and the presence (green) or absence (pink) of TM domains and/or signal peptides. **C.** Table of genes from (A) linked to the pt2 cluster, showing the PlasmoDB annotation and the presence (green) or absence (pink) of a TM domain and/or signal peptides. **D.** Bar plot depicting the total percentage of disordered residues (Y-axis) in the predicted protein structures of the highly expressed, non-annotated genes (X-axis). The color scale indicates the number of disordered segments present in each protein. The 3D structures were analyzed using the CSpritz server.

## Discussion

The transcriptional regulation of *P. falciparum* development in the mosquito midgut remains understudied, with the availability of only a single-cell atlas of *P. falciparum* mosquito stages that includes mature ookinetes (Real et al., 2021). Using scRNA-seq, our data defines the genetics underlying the differentiation of a female gamete to an ookinete. We characterized five transcriptionally unique *P. falciparum* cell states as the parasite develops in the midgut lumen *An. gambiae*, its natural mosquito vetor. We also defined three distinct molecular signatures that orchestrate cell type transitions in the mosquito midgut. Genes related to DNA replication and metabolic processes were identified in early zygotes; reproduction, localization, and motility in intermediate stages, while early to mid-ookinetes express genes related to entry into its host and to down-regulation of metabolic processes. *Plasmodium* development in the mosquito involves a series of DNA replications and meiotic divisions. After fertilization, the zygote undergoes a round of DNA replication that increases ploidy to 4N, followed by a conventional two-step meiotic division (Janse et al., 1986). This is later followed by endomitotic division in the oocyst to produce the sporozoites. We observed the differential expression of genes involved in DNA replication and repair, as well as in meiosis, in three distinct expression programs; early, mid and late midgut development. The early program includes genes like NEK2, NEK4 and MISFIT that has been shown to regulate DNA replication in preparation for meiosis (Guttery et al., 2012). The mid developmental program includes genes like SMC1, CAPD3 and RAD23, involved in chromosomal segregation and DNA repair during meiosis (Hirano, 2016; Kalitsis et al., 2017; Madura & Prakash, 1990). The late program includes proteins expressed exclusively in ookinetes like SMC5, and RAD5 involved in DNA replication and repair before and during mitosis (Hirano, 2016; Ortiz-Bazán et al., 2014). Of interest, PlasMei2, which in *P. yoelii* have been shown to be expressed exclusively in the liver stage where it controls chromosome segregation (Dankwa et al., 2016), was detected in the late program possibly indicating differences in expression with *P. yoelii*. Thus, together with other late program genes, PlasMei2 may have a role during the mitosis of the oocyst sporogony. Differential gene expression analysis determined a single cellular lineage for isolated cells, most of which aligned with the time points of isolation, although a number of individual cells showed temporal transcriptional dynamics aligning with earlier or later time points.

In *P. falciparum*, most transcription factors belong to the Apetala2 (AP2) family which consists of 27 genes. Some of these transcription factors are named according to the life cycle stage where they are expressed and exert their function, for example ApiAp2-g and its role in gametocyte differentiation (Ralph & Cortés, 2019). The progression through the life cycle in the mosquito midgut is dependent on the temporal regulation of ApiAP2 transcription factors, as indicated by the differential expression in pseudotime clusters. With the exception of AP2-O4, all AP2-O genes showed expression in our dataset, and the high resolution provided by single-cell transcriptomics, allowed the definition of the timing of AP2-O gene expression through the studied time course. Moreover, additional ApiAP2 genes appear to be upregulated in a significant proportion of cells (>50%). Two ApiAP2 genes were found in C2 (PF3D7_1239200 and PF3D7_1342900) and four genes in C4 (PF3D7_0420300, PF3D7_0934400, PF3D7_0802100 and PF3D7_1139300), none of which had previously been linked to a developmental stage. In agreement with our data, PF3D7_0934400 was shown to be highly expressed among ookinetes in the single-cell data by Real and colleagues (Real et al., 2021). Also, PF3D7_0622900: AP2Tel, which is significantly upregulated in C4 in our dataset, showed low to intermediate expression among ookinetes in the dataset of Real and colleagues and high expression among a portion of the oocyst-sporozoites. This could be indicative of its involvement in regulating DNA replication genes in oocysts. AP2Tel has been described to be expressed throughout the asexually replicating blood stage in *P. falciparum,* where it was shown to bind directly to telomeric repeats of all 14 chromosomes (Sierra-Miranda et al., 2017). In addition, AP2Tel (AP2-SP3) has been shown to have a role in sporozoite release from oocysts in *P. berghei* (Jeninga et al., 2019; Modrzynska et al., 2017).

The timing of ApiAP2-O expression coincides with the down regulation of three genes: P25, H2B and CPR. These genes are known to be i) a zygote surface protein (Dijk et al., 2010), ii) a component of the nucleosome known for packaging chromatin (Bennett et al., 1995), and interestingly iii), an antimalarial resistance-associated gene in sporozoites (Fan et al., 2011), respectively. The timing of expression of these genes raises the intriguing question of whether AP2-O may be involved in negative regulation of their expression. The involvement of AP2 transcription factors in the negative regulation of genes occurs in *Toxoplasma gondii* (Radke et al., 2018), but has only been speculated on for *Plasmodium spp*. In later time points, AP2-O upregulates an array of genes, among them SOAP and WARP, both important for ookinete stages and important in invasion (Kaneko et al., 2015). Taken together, our results highlight the coordinated expression of ApiAP2 transcription factors during the developmental trajectory of *P. falciparum* in the mosquito midgut and beyond.

Since parasite numbers decline precipitously during mosquito midgut development, we also sought to elucidate the genetic program involved in this critical bottleneck of the *P. falciparum* life cycle. The parasite is inflicted by oxidative stress produced by; free heme resulting from hemoglobin digestion, ROS, toxic molecules produced by the midgut microbiome, human-derived immune and inflammatory molecules, and immune cells ingested with the blood meal (Smith et al., 2014). Subsequently, once the mature ookinete traverses the midgut epithelium, the mosquito complement immune response is activated, leading to increased susceptibility to nitric oxide toxicity (Oliveira et al., 2012). All of these factors contribute to the reduction of parasite numbers seeing during the major lifecycle bottleneck in the mosquito midgut (Griffin et al., 2010; Smith et al., 2014). Hence, we investigated the expression of genes involved in the response against ROS and Reactive Nitrogen Species (RNS), cellular stress and cell death. We found a temporal induction of known stress-related genes indicating that different stressors from the mosquito midgut environment are associated with specific stress responses from the parasite. For example, the redox genes nPrx, HSP90 and dTCTP were highly expressed in the intermediate stages (C1, C0) likely to facilitate survival within the harsh midgut conditions. Previous studies have reported the induction of similar genes, like 2-Cys peroxiredoxin and peroxiredoxin-1, due to the environment of the mosquito midgut and their role in protecting the parasite midgut stages from oxidative stress (Peterson & Luckhart, 2006; Turturice et al., 2013).

Previous studies have used I-TASSER (Yang & Zhang, 2015) to elucidate the structure and function of proteins of interest, including *in silico* experimentation for finding novel antimalarials (Pandey et al., 2018). We used I-TASSER to predict protein function of a set of highly expressed, non-annotated genes linked to the mid and late developmental points. Using 3D model predictions, we selected 17 non-annotated and highly expressed genes with a predicted TM domain and/or a signal peptide, indicating their possible membrane localization. These proteins were further indicated to be intrinsically disordered proteins (IDPs), proteins largely disordered in their structure and reported to be highly immunogenic (Guy et al., 2015), a trait essential for antibody-based therapies.

The *P. falciparum* midgut stages represent a tenuous point in the parasite’s life cycle given the large decline in population size and genetic diversity. Such small numbers render emergence of parasites resistant to therapeutics far less likely, as compared to human stages. These properties make the *Plasmodium* mosquito stages a highly strategic point for malaria intervention. Drugs which target this stage will be highly specific against the parasite and can be administered to cure mosquitos and indirectly prevent human infection (Paton et al., 2019). Further, human antibodies are potent, remain viable within the mosquito midgut and are becoming affordable for use in low income settings, indicating that transmission-blocking vaccines targeting the transmission stages of the parasite are possible and merit further exploration (de Jong et al., 2021; Kundu et al., 2018; Sauerwein & Bousema, 2015). Overall, our data provides an important resource for exploring the hitherto largely undescribed transcriptome of the mosquito midgut stages of *Plasmodium falciparum*, and offers untapped potential for the exploration of transmission-blocking therapeutics.

## Materials and Methods

### *In vitro* culture of *P. falciparum* parasites and gametocyte induction

Gametocytes from the *P. falciparum* gametocyte-producing NF54 cell line were induced and cultured as previously described in (Delves et al., 2016). In brief, parasites were cultivated on type O positive erythrocytes at 4% hematocrit in standard culture media containing RPMI supplemented with 10% heat inactivated human serum (Karolinska Hospital blood bank, Stockholm, Sweden) using standard culturing techniques. Cultures were gassed with 96% N2, 1% O2 and 3%CO2 and maintained on a shaking incubator at 37 °C to prevent multiple infections in RBCs. Gametocytes were then induced at 5-6% parasitaemia and maintained, with daily media changes for 14 to 17 days in gametocyte culture medium containing 25 ml of RPMI and 10% heat inactivated human serum. Media were changed on a daily basis after which mature gametocytes were harvested for mosquito infection on days 14 and 17.

### Mosquito infections

Gametocyte cultures were harvested on day 17 and diluted to 1% gametocytemia at 50% hematocrit, and delivered directly into water-jacket glass-membrane feeders connected to a 37°C circulating water bath. Twenty *Anopheles gambiae* females (5-7 day old), maintained at 25-27°C and 80% humidity, were allowed to blood-feed for about 30 minutes, after which any unfed mosquitos were isolated and discarded. Between 15 to 20 mosquito midguts were dissected at time points 2, 4, 8, 12 and 20 hours post infection. Mosquito midguts were collected in 1.5 ml Eppendorf tubes with 500μl of RPMI and homogenized using a micro-tube homogenizer (F65000-0000, SP-Bel-Art) at short intervals for 30 seconds. In addition, one batch of mature gametocytes were treated with aphidicolin (Sigma #A0781-1MG) *in vitro* and cultured for 2h at room temperature. Aphidicolin prevents DNA replication in male gametes rendering them immature and thus blocks female fertilization.

### Staining, imaging, and isolation of single parasites

After midgut homogenization, 2μl of anti-Pfs25 antibody (1 µg/ml) was added to the sample and incubated at RT for 15 minutes. 1 μl of Alexa Fluor 488 goat anti-mouse IgG (H+L) at 2 µg/ml (Life Technologies, cat#A11001) and 0.5 μl of 10 mg/ml Hoechst 33342 was added to the sample and incubated for 15 minutes at RT in the dark. Cells were washed two times with sterile DPBS (Dulbecco’s Phosphate Buffered Saline, ThermoFisher Scientific, cat#A1285801), parasite pellets were subsequently resuspended in 500 μl of DPBS. 100 μl of stained parasites were placed on BSA coated petri dishes to prevent the adhesion of the parasite to the glass bottom of the petri dish. Targeted individual parasites were visualized and imaged using a Leica DMi8 microscope with K5 camera, 4.2 megapixel (Leica, Germany). Targeted cells were collected using capillary-based micro-manipulation with glass capillaries containing an inner diameter of 8µm (Eppendorf, Hamburg, Germany). Capillaries were pre-coated with a sterile 2% BSA solution to prevent the gametes from sticking to the glass capillary surface. Z-stacks taken of targeted single parasites and were then processed using IMARIS software (Version 9.3). Image processing analysis for each time point was set to a standard and was used to measure the mean fluorescence intensity and the volume of the nucleus against different cell types yielded by clustering analysis. Individual parasites were then captured and replaced in 1x DPBS and transferred into 200 μl thin-walled PCR tubes (Corning, NY) containing 3.5 μl of lysis buffer (0.6% Triton-X100) 2U/μl recombinant RNase inhibitor, 1 μl oligo-dT (10 μM) and 1 μl dNTP mix (10 mM). All samples were immediately stored at -80°C after isolation.

### cDNA synthesis and library preparation

cDNA libraries of single parasites were generated using a modified version of the Smart-seq2 protocol (Picelli et al., 2014). In short, cDNA synthesis was performed using *P. falciparum* optimized primers (Real et al., 2021) and PCR amplification was carried out over 24 cycles. cDNA products were subsequently purified using CA Beads (Sigma, Cat. Nr. 81260) for size selection using 8.8% PEG6000 to exclude primer-dimers and non-specific amplicons with sizes less than 150bp. Combinatorial indexing via tagmentation was carried out in 96-well plates using 200pg (measured in a Qubit) of amplified cDNA, for a final volume of 10 μl/well. cDNA fragmentation using Tn5 transposase was carried out for 20 minutes on ice using the Illumina Nextra XT DNA sample preparation kit. Ligation and amplification of adaptors was carried out over 15 cycles in a final volume of 25 μl/well. Primer indices were used in the reaction from illumina (Nextera index primers-i7 and i5 cat# FC-131-1001). Tagmented and barcoded amplicons were then purified using CA Beads for size selection. Quality control and fragment size distribution of the cDNA libraries were performed on a BioAnalyzer with the Agilent high sensitivity DNA chip cat# 5067-4626. Concentrations of each sample of cDNA libraries were measured on a Picogreen 96-well plate NucleoScan fluorometer using a high sensitivity dsDNA (HS assay kit, cat# Q32851). To perform library dilutions, the average fragment sizes of all cDNA libraries were measured for a final concentration of 2nM in each sample. Finally, cDNA libraries were pooled and sequenced using Illumina NextSeq with 75 bp paired-end reads.

### Computational analysis

#### scRNA-seq raw data mapping and feature counts

Raw scRNA-seq reads were processed and trimmed for quality and adapter content using FastQC (Version 0.11.5) (Andrews S. 2012) and Trimmomatic (Version 0.36) (M. Bolger 2014) respectively. All quality processed reads were mapped to Human (GCF_000001405.38_GRCh38.p12_genomic.fna), mosquito vector (GCF_000005575.2_AgamP3_genomic.fna) and *P. falciparum* parasite (GCF_000002765.4_ASM276v2_genomic.fna) using FastQ_Screen, and the reads mapping to parasite but not human and vector were retained. All orphan unpaired reads were discarded and remaining paired reads were mapped to *P. falciparum* genome downloaded from PlasmoDB (PlasmoDB-39_Pfalciparum3D7_Genome.fasta) using STAR (Version 2.5.3a) (Dobin A. 2013). Read quantification was performed using HTseq-count (Anders S. 2015) (parameters:-t exon -i gene_id -r pos -m intersection-nonempty) and custom bash script were used to generate gene count matrix.

### Quality control and normalization

For estimation of good quality cells to perform downstream analysis we used the Seurat package (v 4.0.3) on the raw features count matrix for an overview of the distribution of the number of reads and genes detected per cell within each time point. We set a cutoff to filter out cells that had fewer than 600 genes and 2500 reads resulting in 125 cells that were included for the downstream analysis. We then employed a global-scaling “LogNormalize” to the feature expression measurements for each single cell from the total expression multiplied by a scale factor (10,000) with the normalized values stored in the Seurat object.

### Identification of highly variable features

In order to delineate cell-to-cell variation across the dataset and highlight the biological signal of highly expressed genes, we used the feature selection method “vst” implemented in the Seurat package (FindVariableFeatures) to find variable features across the single cells’ transcriptomes. The top 2000 variable genes were selected for the downstream analysis.

### Dimensionality reduction, cell clustering and projection

To dissect cellular heterogeneity during the development of *P. falciparum* zygotes, we first scaled the expression so that the variance across cells had an equal weight of 1 in the downstream analysis. We then performed linear dimensionality reduction (principal component analysis – PCA) using the top highly variable genes (HVGs) on the normalized-scaled transcriptome data to characterize the variation across the dataset. To estimate the number of PCs indicating a true signal of transcriptional variation, we visualized both the cells and features using the DimHeatmap function on the first 20 PCs and then performed the JackStraw test for the P-value estimation. Next, we used the first 10 PCs (most PCs showing the signal variation) to delineate the transcriptome into cell type communities by running kNN graph-based clustering and applying modularity optimization with the Louvain algorithm to relatively group the cells with a resolution = 0.8. Different cell types’ transcriptomes were then visualized using UMAP non-linear dimensionality reduction. Detailed steps of the analysis are explained in the data and code availability section.

### Differential gene expression analysis

To define markers distinguishing each cluster generated by the Louvain algorithm, we set a minimum percentage of cells expressing certain features to be detected at 25%. Finding differentially expressed genes across the cluster was set with Logfc.threshold of 0.25 using the “Wilcoxon rank sum test” with a minimum of 25% cells for cluster specific markers comparing all remaining cells reporting positively and negatively regulated markers to be used in GSEA (Gene set enrichment analysis) for particular biological pathway analyses. The differentially expressed gene dataset is presented in Supplementary Table 3.

### Cell lineage and Pseudotime analysis

Louvain clusters were ordered along developmental trajectory using Slingshot (v1.8.0) (Street et al., 2018). Non-fertilized female gametocytes (NFFGs) were excluded and the feature expression matrix was normalized and scaled to estimate the total mRNA abundance with the aid of the Seurat object tool along with metadata previously generated from Louvain clustering. To restructure cell lineage and develop a pseudotime inference for the purpose of uncovering the global mRNA structure of the single-cell dataset, we used the MST (minimum spanning tree) algorithm and fit our cells simultaneously against a “principal curve” to establish mRNA distribution in an unsupervised manner. Applying the MST recovers a single lineage of developmental trajectory composed of three global cell communities (Pseudotime regulatory modules). Based on differentially expressed features recovered using the FindAllmarkers function, specific cluster markers were identified and (C3) were assigned as rooting cells or the lineage starting point (Initial cells) of the developmental trajectory while the pseudotime values were estimated simultaneously for each single cell ordered along the fitting curve. To visualize the pseudotime values specifically for the Seurat clustering and the isolated time points, we overlaid the single cells’ metadata to the Slingshot lineage to have a better understanding of the relationship between the clusters and the isolated time points (Figure 2B). To find genes that change their expression over the course of the development, we used the Tradeseq package 1.4.0 (Van den Berge et al., 2020) to calculate the relationship between gene expression and pseudotime. In brief, we used a general additive model (GAM) to model the relationship between genes and conduct an association test to estimate the *p*-value of genes significantly expressed over time (Figure 2C and Supplementary Table 4). We summarized snapshot transcripts of the global structure lineage and notable GO terms from PlasmoDB.org were used to distinguish pseudotime gene co-expression modules (pt0-pt2). To assess genes transcriptionally regulated during the course of development with significant variation in their expression, we employed the global clustering structure forming slingshot single lineage.

### Motif identification for ApiAP2 regulators

We explored known ApiAP2 target sequence motifs using a Plasmodb search on transcriptionally regulated genes generated from pseudotime analysis (differentially expressed genes along the pseudotime) which returned only the genes with a specific known binding mix-base motif within a 1 kb upstream region of the starting codon for each ApiAP2 transcription factor target genes. Next, we visualized the target gene dynamics of expression in relation to the specific transcription factor showing the regulation of target genes over the course of pseudotime (Kaneko et al., 2015).

### Biological pathways and gene set enrichment analysis (GSEA)

To track tests for top functional class enrichment among the global clusters building the pseudotime lineage, we used conservative markers generated for each cluster on PlasmoDB GO analysis tool to conclude the enriched ontology terms as previously mentioned. The GSE analysis was performed on differentially expressed genes over the pseudotime as input in cluster profiler v 3.18.1 and ggupset package v 0.3.1 with a p-value cut off = 0.05, minGSSsize = 3, maxGSSize=800 and scoreType=” pos’’ to estimate for biological process ontology changes over the pseudotime lineage and developmental progress (Supplementary Figure 3A and Supplementary table 1). The top 20 biological processes were visualized using the Clusterprofiler package (v 3.18.1) dot plot function.

### Protein prediction analyses

The amino acid sequences of significantly upregulated, non-annotated genes of *P. falciparum* 3D7 were retrieved from the PlasmoDB website and processed for their primary structures and physico-chemical properties. Briefly, Expasýs ProtParam tool (Colovos & Yeates, 1993) was utilized to calculate the physico-chemical characteristics and the secondary structural properties including α helix, 310 helix, Pi helix, Beta Bridge, Extended strand, Bend region, Beta turns, Random coil, Ambiguous states and other states using MLRC/SOPMA (Guermeur et al., 1999) I-TASSER (Yang & Zhang, 2015) was used for the structure-based functional annotations. Predicted proteins with high confidence prediction annotations were then examined for the presence of signal-peptide or TM domains. Alpha-Fold (protein structure database) was utilized to retrieve the .pdb files of the predicted protein structures (Jumper et al., 2021). I-Tasser Modrefiner was used for structure refinement (Xu & Zhang, 2011) based on atomic-level energy minimization. The accuracy and stereochemical features of the predicted models were then calculated with PROCHECK on the PDBsum server (Laskowski et al., 1996) using “protein structure and model assessment tools”. Three putative membrane protein 3D models were then subjected to druggability assessment using DogSitescorer server for druggable pocket predictions (Supplementary Figure 7).

### Identification of intrinsically disordered proteins (IDP)

We computationally characterized the 3D homology structures of protein models showing a lack of well-characterized protein segments (IDP) using CSpritz server for accurate detection of protein disorder (Walsh et al., 2011) in combination with comparative dynamic simulations with python-based docking libraries (py3Dmol, openbabel and nglview). Briefly, .pdb files retrieved from Alpha-fold and imported in Jupyter notebook with perquisites libraries imported and compared with well-structurally annotated proteins. We then identified the characteristics of bulky hydrophobic amino acids sequences with high net charges promoting disorder in form of extended loops regions coupled with folding and binding compared to the core structure resulting in instability and irregularity of the secondary structure (van der Lee et al., 2014).

### Data integration of 24-hour Ookinetes (Day 1) Real et al 2021

Expression matrices and supporting metadata files were downloaded from https://zenodo.org/record/4719664/. Single-cell data were subset according to the target stage (Day 1 Ookinete). Variable genes were intersected between the two datasets using Scanpy (v1.5.0) (Wolf FA. 2018). We then used the concatenate and ingest function to integrate data annotations and labels and corrected for batch effect using BBKNN integrated in scanpy workflow.

### Data and code availability

In-house bash, R code scripts and data that were implemented in this study are available on GitHub [https://github.com/ANKARKLEVLAB/Single-cell-P.falciparum-midgut]. Expression matrices and meta data are available via [https://zenodo.org/deposit/4683823] and the data is also searchable via [https://mubasher-mohammed.shinyapps.io/shinyapp/].

## Acknowledgment

We would like to thank The Microbial Single Cell Genomics (MSCG) facility, SciLifeLab, for assistance with the single-cell sorting and optimization of molecular workflows and library preparations and the National Genomics Infrastructure (NGI) for Computing (SNIC) for data handling and preprocessing of the scRNA-seq raw data for this study, partially funded by the Swedish Research council through grants agreement 2018-05973 to J.A. The computations were performed using resources provided by SNIC through Uppsala Multidisciplinary Center for Advanced Computational Science (UPPMAX) under Project SNIC 2020/16-146 and SNIC 2021/22-492.

This work was supported by grants from the Swedish Society for Medical Research (SSMF), The Swedish Research Council (VR-NT), The Jeansson Foundation and SciLifeLab to J.A.; The NIH Distinguished Scholars Program and the Intramural Research Program of the Division of Intramural Research AI001250-01, NIAID, National Institutes of Health; the Strategic Research Area (SFO) program of the Swedish Research Council (VR) through Stockholm University to V.S and M.R.F; the Swedish Research Council (VR) #2021-06602 to J.H.; and NIH grant R01AI031478 to M.J.L..

## Authors contributions

**J.A.** and **J.V.R.** conceived the study; **J.A., J.V.R., M.R.F.** and **M.J.L.** supervised the project; **M.M., J.V.R.** and **J.A.** performed cell culture, infections, dissections, staining, cell imaging and micromanipulation for cell isolation; **N.E.** provided technical support and assistance with mosquito culturing and infections; **M.M.** performed Smart-seq2, library preparations; **M.M.** and **I.B.** performed sequencing; **V.S.** performed scRNA-seq data preprocessing and quality control; **M.M.** performed computational analysis, data integration and interactive web visualization; **B.B.** and **A.J.** performed the iTasser analyses. **J.H. J.A. and M.R.F.** supervised the computational analysis; **J.V.R., M.K., T.S.** performed the imaging analysis; **A.D.** conceptualized and coordinated data visualization, **J.A., M.M., AD** wrote the manuscript with help of **J.V.R., M.J.L.** and **J.H**, all authors edited and provided critical input.

## Additional information

**Supplementary information is available for this article.**

Correspondence and requests for materials should be addressed to

---

**Supplementary Fig 1.**
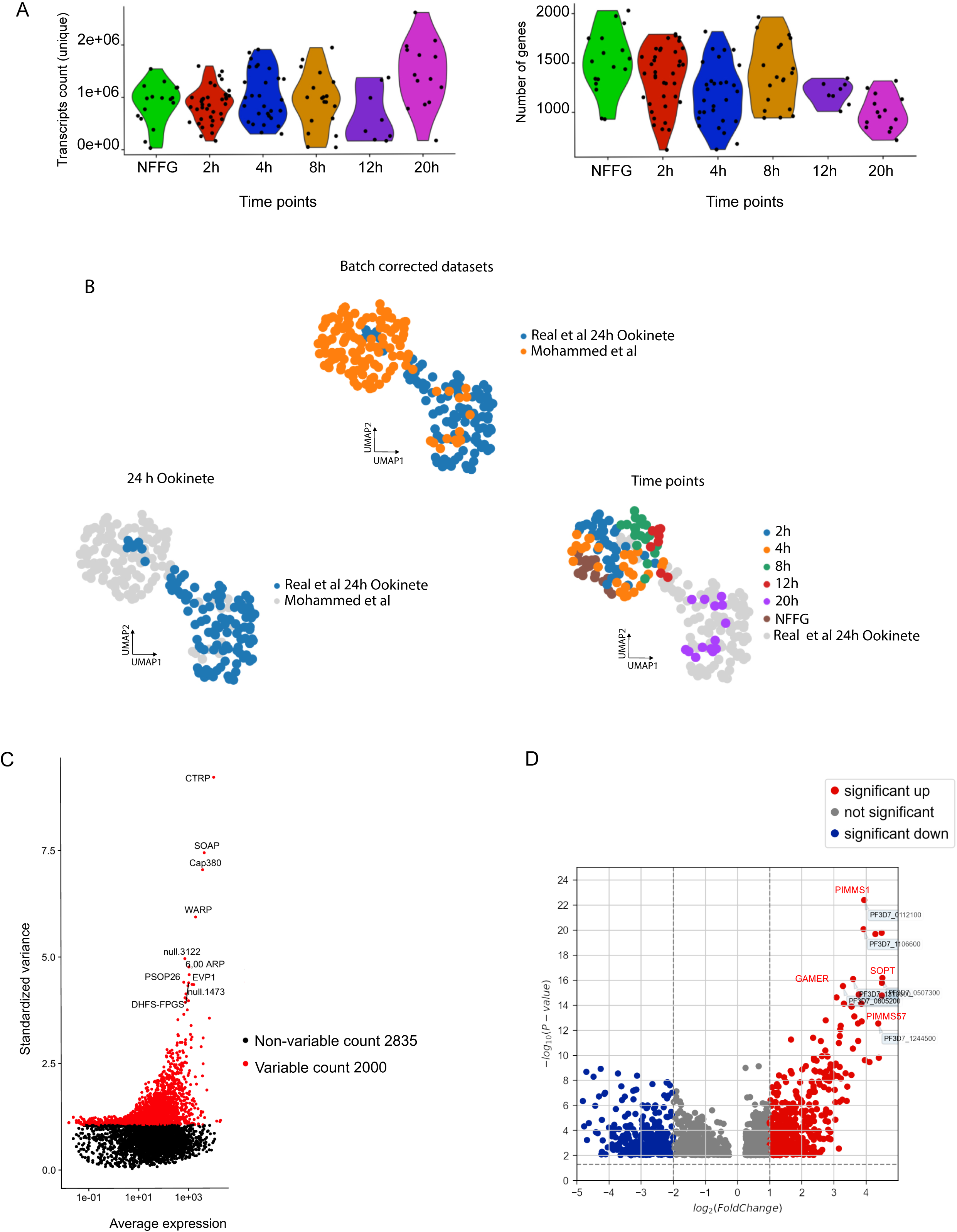
Quality assessment of Smart-seq2 data and data integration with Real *et al*. 2021. **A**. Violin plots showing the number of unique transcripts detected per cell (Left panel) and the number of genes detected per cell (Right panel) across the six collection timepoints. **B**. Data integration of Real *et al*. 2021 day 1 (24h) single-cell transcriptomes with time points determined in this work non-fertilized female gametes (NFFG) to 20 h single-cell transcriptomes. Upper panel: shows integrated datasets. Bottom panel left: integrated datasets highlighting (blue) 24h ookinetes from Real *et al*. . Bottom panel right: integrated datasets highlighting the isolated time points determined in this work. After batch correction, more single cells from this work represent early development of *P. falciparum* in mosquito midgut stages. **C**. Volcano plot depicting the most highly variable genes, with the top 10 genes labelled. A total of 2,835 features were identified as non-variable while 2,000 features were variable and selected for downstream analysis. **D**. Volcano plot showing differentially expressed genes, based on Seurat cluster comparison, color-coded as red for significantly up-regulated, blue for significantly down-regulated, or Grey for not significant. The x-axis represents the fold change while the y-axis represents the *P*-value. A total of 1,576 differentially expressed genes were identified

**Supplementary Fig 2.**
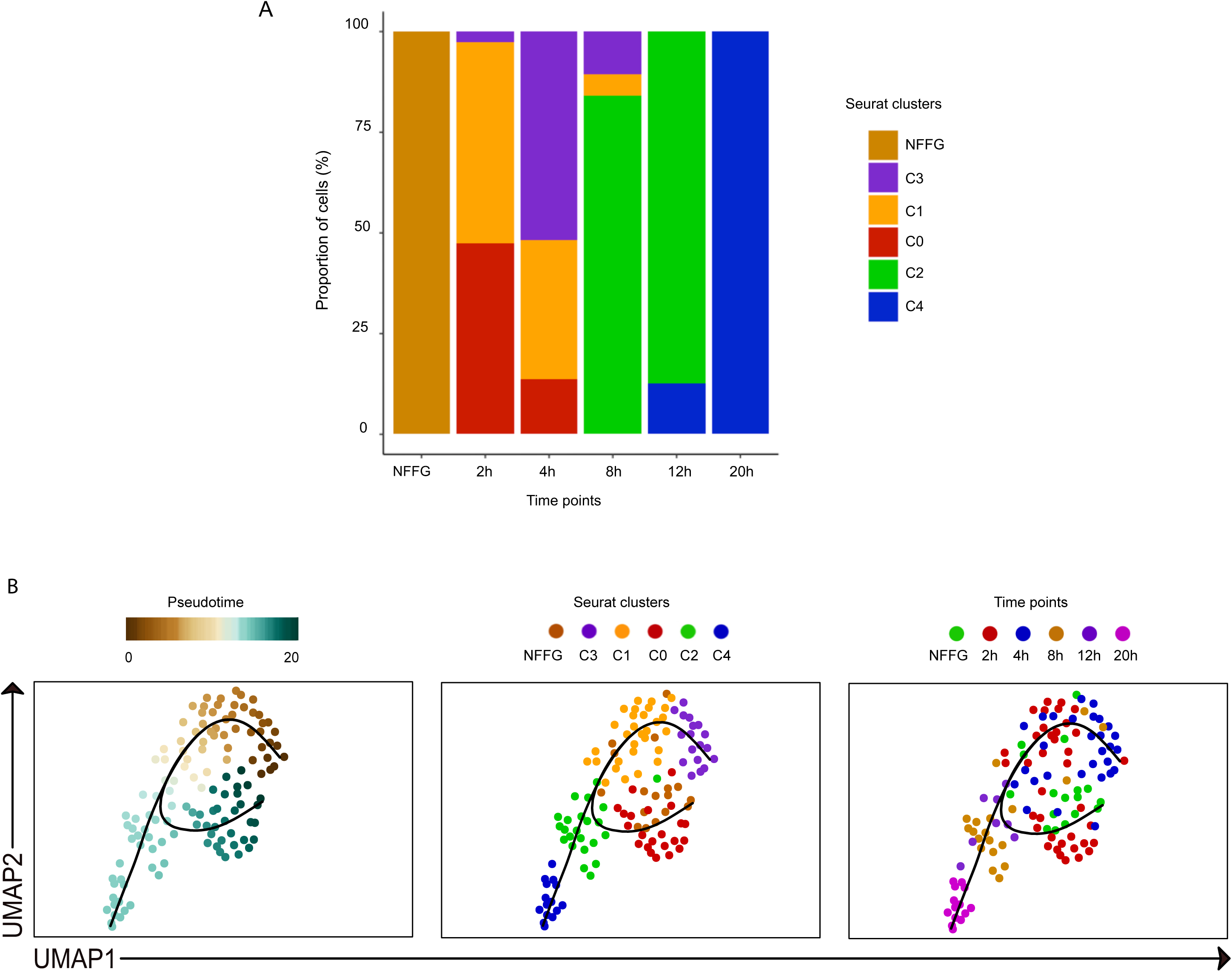
Comparison of the Slingshot trajectory analysis in the presence or absence of the non-fertilized female gametes (NFFG) **A**. Bar plot showing the ratio of Seurat clusters identified within each collection time point. **B.** The UMAPs indicate a modified trend of the Slingshot pseudotime trajectory when NFFGs are included in the analysis as compared with the trajectories shown in Figure 2A, where the inclusion of the single-cell transcriptomes from NFFGs appears to distort the global lineage reconstruction. Left panel: The alternative pseudotime trajectory where the color coding represents the cells predicted development along the pseudotime axis. Middle panel: The pseudotime developmental trajectory overlaid with the assigned Seurat clusters and where NFFGs are separately color coded in brown. Right panel: The pseudotime developmental trajectory overlaid with the cell isolation timepoints and including NFFG.

**Supplementary Fig 3.**
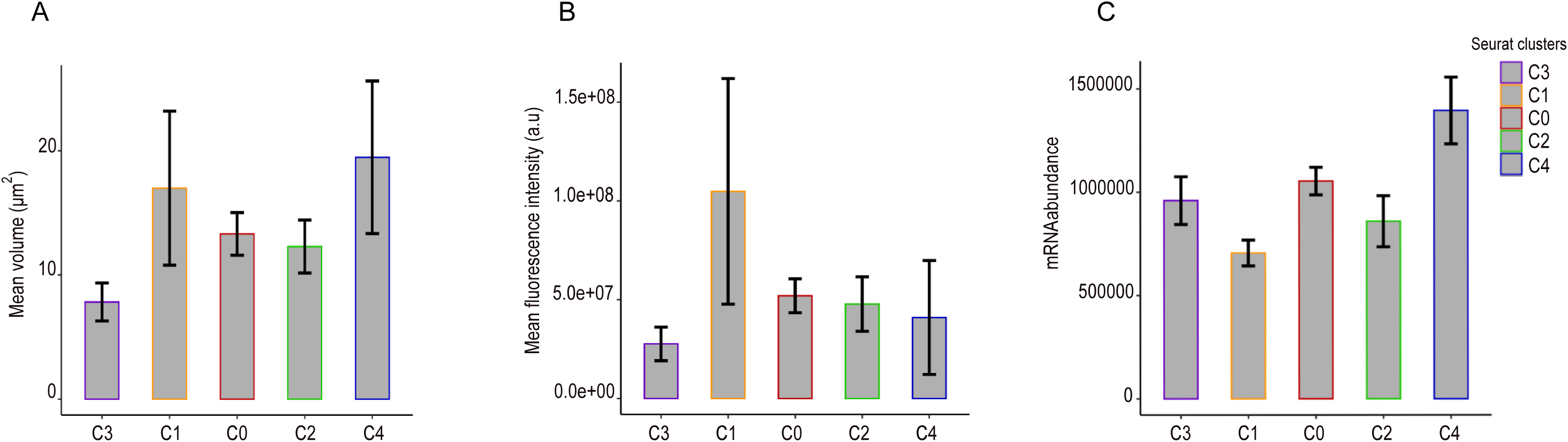
Imaging parameters of ordered Seurat clusters leveraging Pseudotime in characterizing nuclear phenotype features. **A**-**C**. Bar graphs showing the mean volume of the nucleus with standard errors (A), mean fluorescence intensity (B) and mean value of total mRNA abundance (C), among ordered Seurat clusters.

**Supplementary Fig 4.**
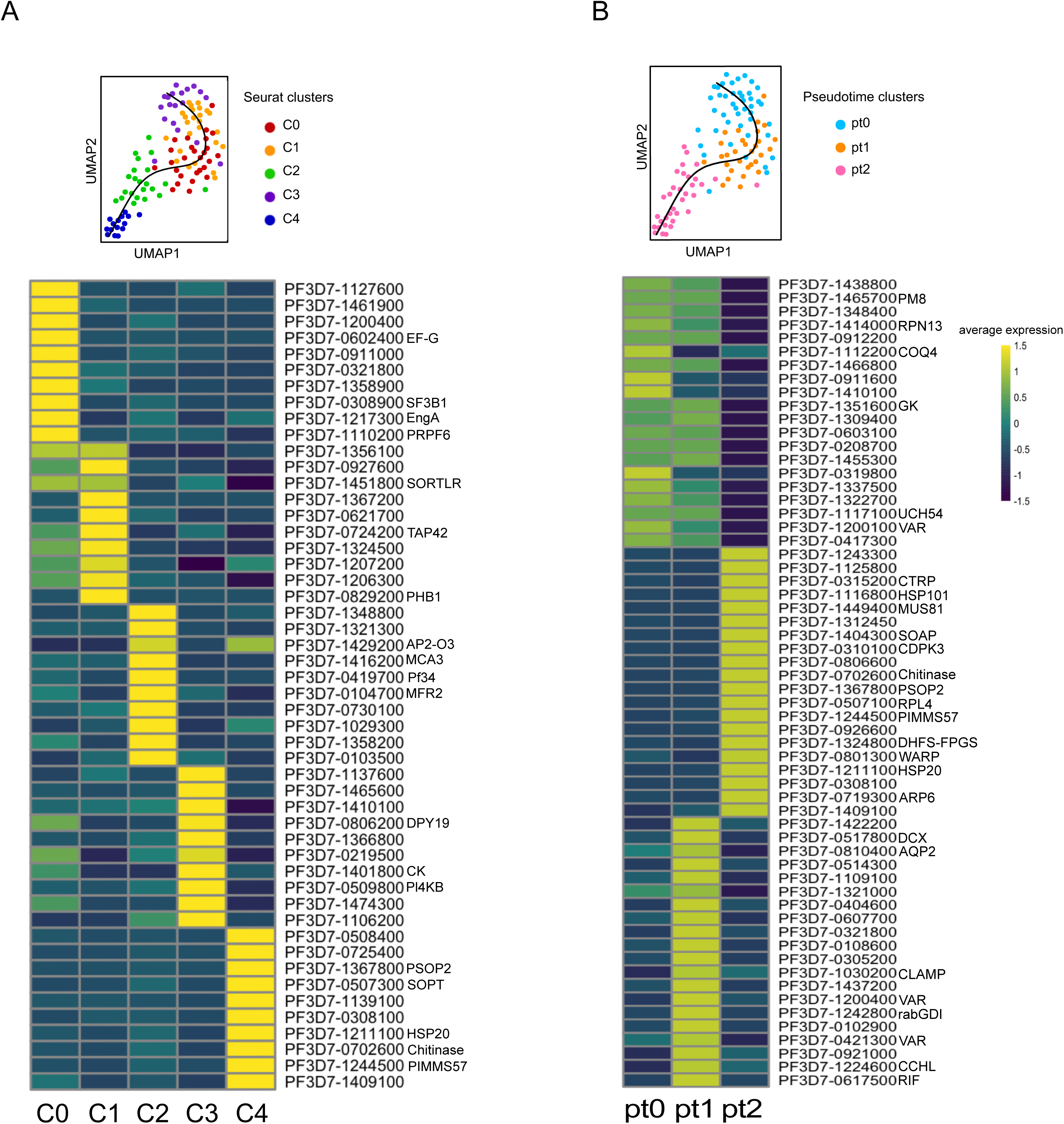
Comparison of top differentially expressed genes between the Seurat and Slingshot clusters along the pseudotime trajectory. **A**. The Slingshot pseudotime developmental trajectory overlaid with the Seurat clusters (top) with a hierarchical clustering heatmap showing the top 10 differentially expressed genes for each individual cluster (bottom). **B**. Slingshot pseudotime developmental trajectory overlaid with the Slingshot pseudotime clusters (pt0, pt1, pt2) (top) with a hierarchical clustering heatmap of the top 20 differentially expressed genes for each individual cluster (bottom). The Wilcoxon rank test was used in the differential gene expression analysis (for both A and B) of genes significantly expressed in at least 25% of cells and logFC threshold of 0.25.

**Supplementary Fig 5.**
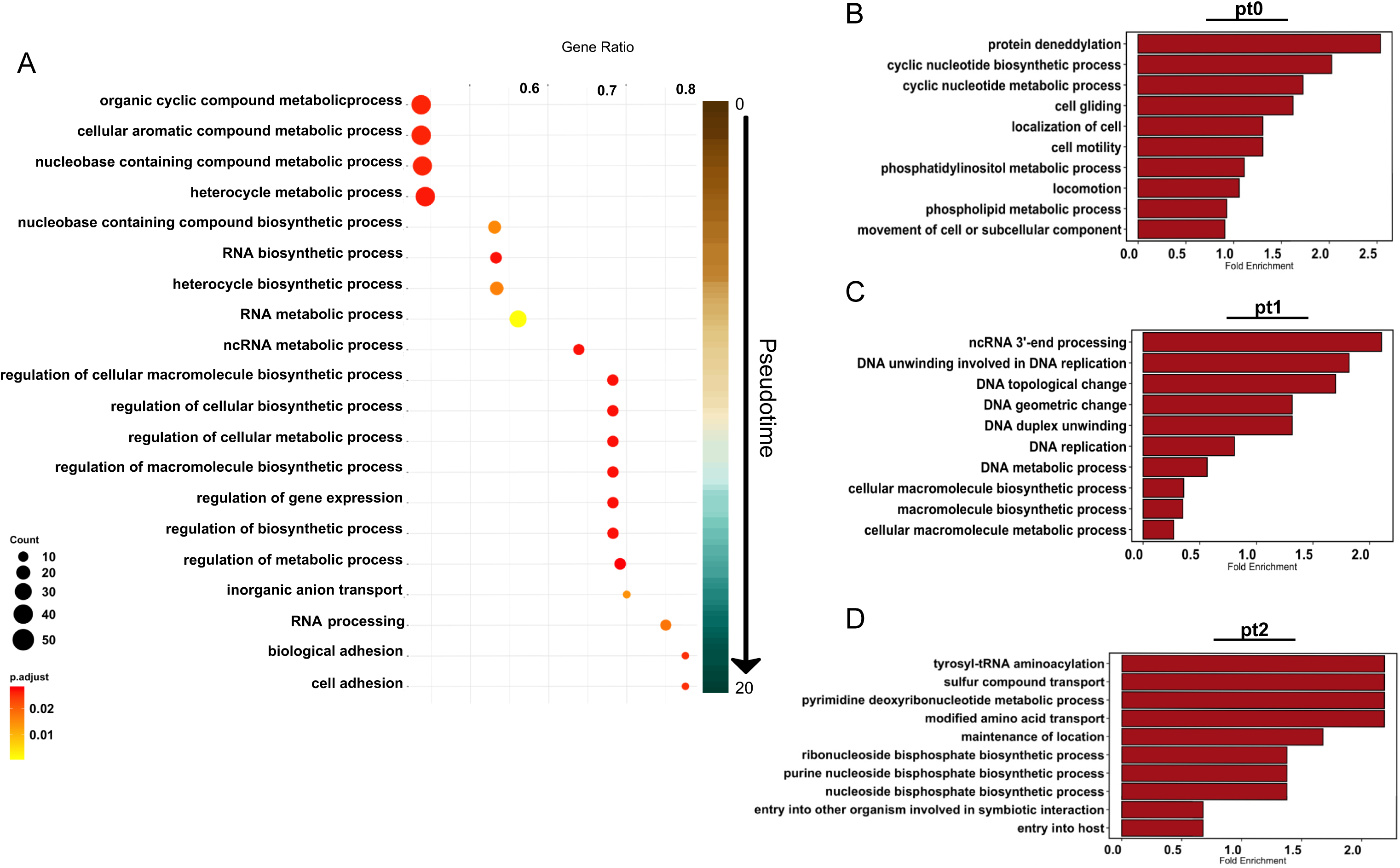
Gene ontology enrichment by Slingshot clustering. **A**. Dot plot of a gene set enrichment analysis from the top differentially expressed genes over the pseudotime. Genes of interest were grouped according to the top 20 GO terms identified in the three pseudotime clusters and statistically analyzed by gene ratio with more significantly (red) and less significantly (yellow) expressed genes, color coded and organized by gene count (dot size). The X-axis indicates gene ratio, and the Y-axis indicates GO terms. **B**-**D**. Bar plots of top GO terms showing the over-represented GO terms for three (pt0, pt1, pt2) cell communities determined using Slingshot pseudotime curve reconstruction. Top 10 GO terms were selected per cluster.

**Supplementary Fig 6.**
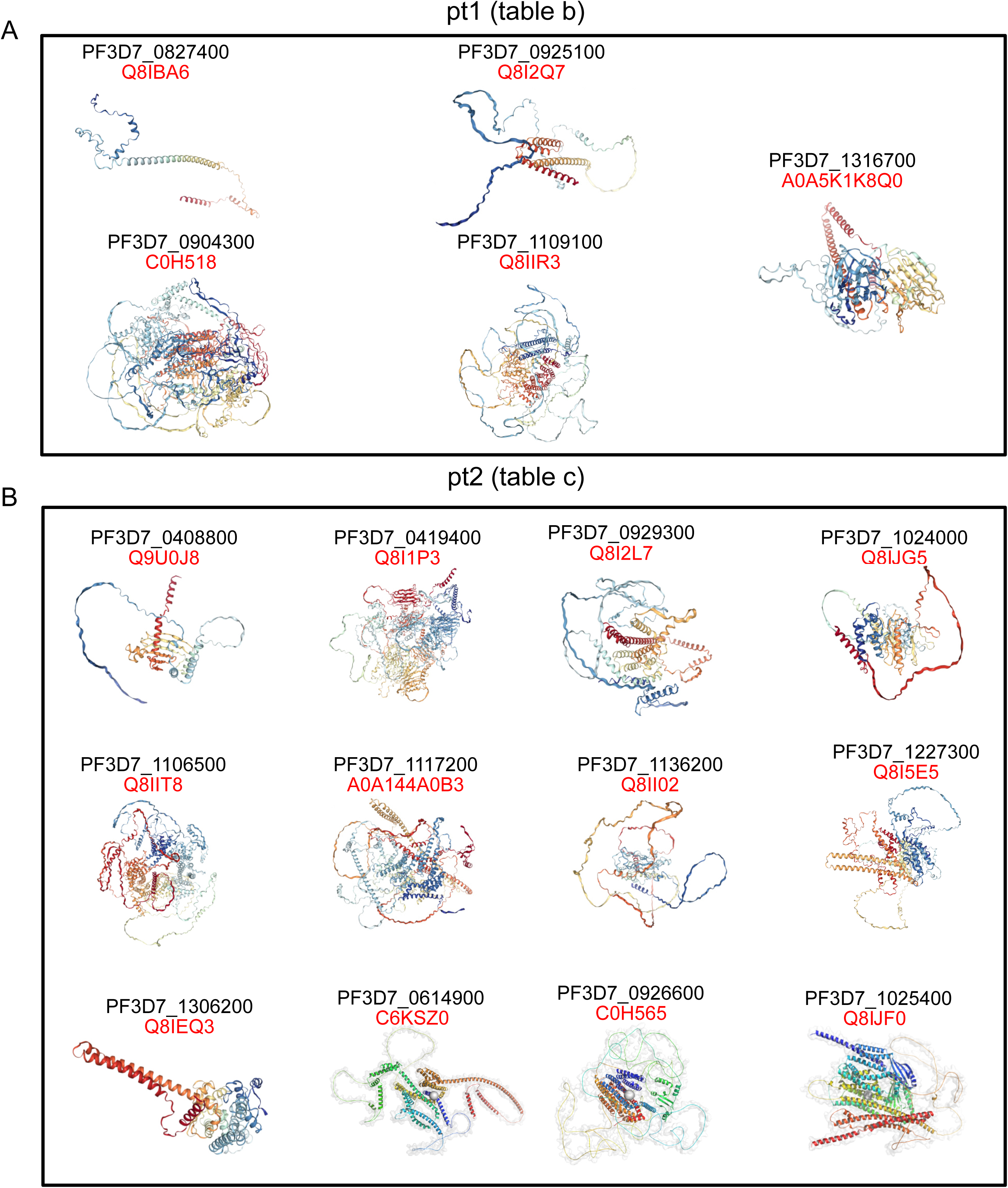
3D models of proteins of the highly expressed genes from pt1 and pt2. Visualization of 3D structure models of intrinsically disordered proteins showing the cartoon topology. The molecular structures were retrieved from the Alpha-fold database and subjected to structural validation using the CSpritz and molecular dynamics simulation tools. **A**. Predicted 3D structures of the non-annotated genes from the pt1 cluster, including their PlasmoDB accession number and UniProt code. **B**. Predicted 3D structures of the non-annotated genes from the pt2 cluster, including their PlasmoDB accession number and UniProt code.

**Supplementary Fig 7.**
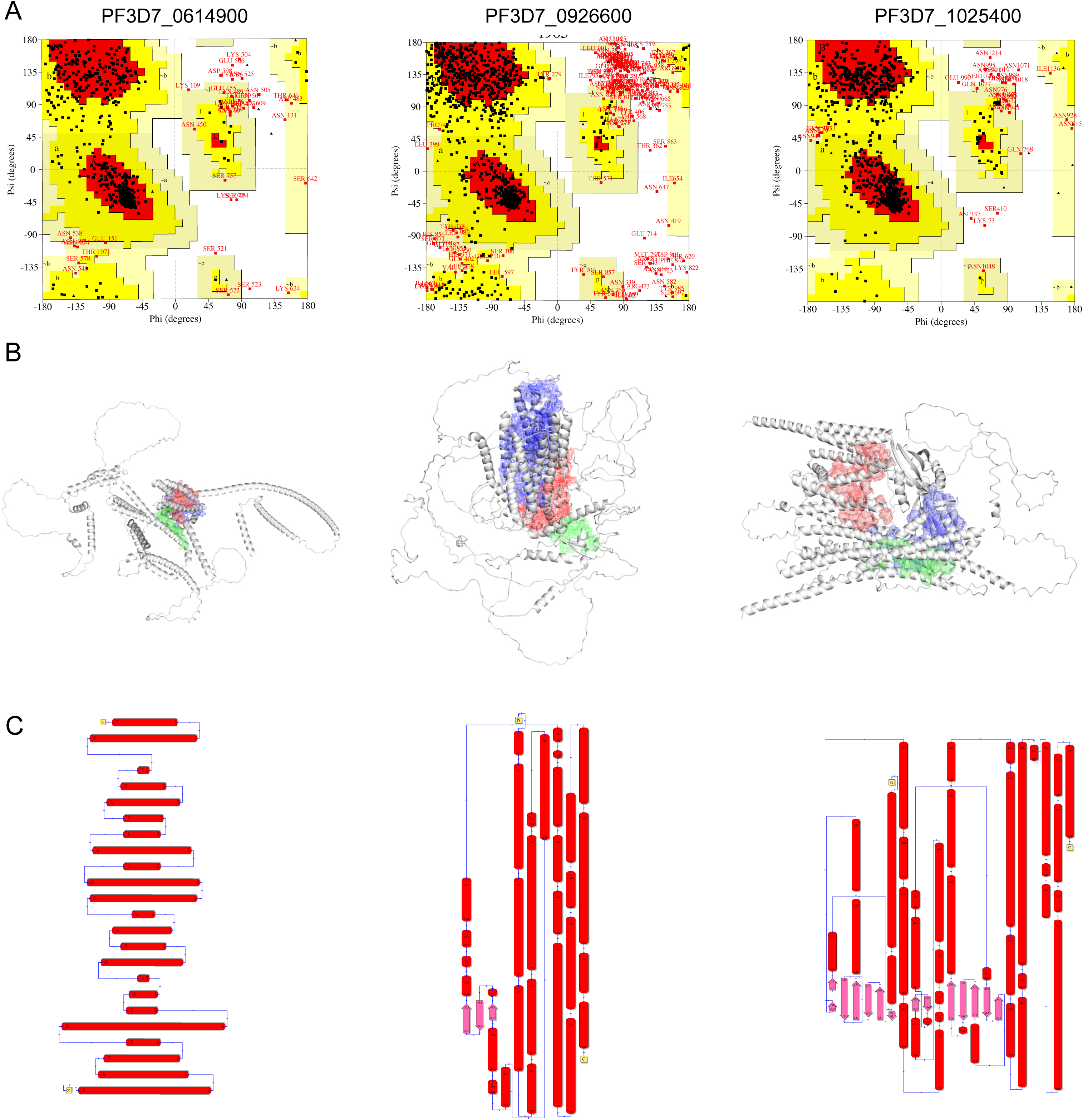
Protein structure and topology to predict the druggability of three putative membrane proteins. **A**. Ramachandran plot analysis for the three predicted proteins described as putative membrane proteins in PlasmoDB. Red regions indicate the most favored amino acid residues, yellow regions indicate allowed regions, light yellow generously allowed regions and white indicating disallowed region. Phi and Psi scales indicate the predicted torsion angles. **B**. Graphical structures and surface topography of the three predicted protein models depicting the largest three binding pockets predicted by DoGSitescorer and simulated using pyMol. The binding pockets are colored blue, red and green. **C**. Topographies for the three predicted membrane protein models from the N terminus to the C terminus, the amino acid residue positions are visualized within the peptide chain.

